# HIV-1 Vpr Induces Degradation of Nucleolar Protein CCDC137 as a Consequence of Cell Cycle Arrest

**DOI:** 10.1101/2021.03.16.435666

**Authors:** Laura Martins, Ana Beatriz DePaula-Silva, Vicente Planelles

## Abstract

Expression of HIV-1 accessory proteins Vif and Vpr results in G2/M cell cycle arrest by hijacking the host ubiquitin-proteasome system. Vif directs cell cycle arrest by targeting protein phosphatase 2, regulatory subunit B alpha (PP2AB56) for degradation. However, the ubiquitination target(s) of Vpr that is directly responsible for G2/M arrest has remained elusive. Recently, Vpr directed degradation of nucleolar protein coiled-coil domain containing 137 (CCDC137), also known as retinoic acid resistance factor (RaRF), has been implicated as the proximal event leading to G2/M cell cycle arrest. In this study we aimed to further investigate this finding. We confirm that CCDC137 is targeted for degradation in the presence of Vpr with a requirement for the CUL4A^DDB1.DCAF1^ E3 ligase complex. However, degradation of CCDC137 is a general consequence, rather than a trigger, of G2/M arrest. Thus, whether induced by Vpr expression or pharmacologically via CDK1 inhibition, G2/M blockade results in degradation of CCDC137. Furthermore, siRNA-mediated depletion of CCDC137 failed to induce G2/M arrest.

## Introduction

Many viral infections in plants and animals result in altered cell cycle progression to benefit host immune system evasion, viral replication, and cell transformation (Davy and Doorbar, 2007; Fan et al., 2018; Qi and Zhang, 2019). Subversion of the host cell-cycle is accomplished through diverse mechanisms mediated by viral proteins or simply through the presence of viral genomes (Davy and Doorbar, 2007). Viral protein-mediated cell cycle arrest is often achieved by modulating gene expression or protein-protein interactions that result in activation or repression, re-localization, or altered stability of host cell cycle regulatory proteins (Davy and Doorbar, 2007; Fan et al., 2018). One common method of virus-mediated host protein destabilization is through manipulation of the ubiquitin-proteasome pathway (Calistri et al., 2014).

Two HIV-1 accessory proteins, viral protein regulatory (Vpr) and viral infectivity factor (Vif), have been shown to induce G2/M cell cycle arrest in CD4 T-cells (He et al., 1995; Jowett et al., 1995; Re et al., 1995; Sakai et al., 2006). Proteomic analysis of HIV-1 infected cells indicates that the presence of Vif significantly alters protein abundance of 15 proteins (Greenwood et al., 2016). In contrast, a similar study identified 1,900 significant protein abundance changes associated with Vpr (Greenwood et al., 2019). Both Vpr and Vif hijack E3 ligase complexes to degrade antiviral targets. Vif uses a cullin 5 (Cul5) E3 ligase complex to degrade several apolipoprotein B editing complex family members (APOBEC3), which are cytidine deaminases, preventing APOBEC3 packaging within newly assembled virions in producer cells and thus preventing hypermutations of viral genomes (reviewed in (Desimmie et al., 2014)), and PP2AB56 for degradation resulting in the induction of G2/M cell cycle arrest ((DeHart et al., 2008; Greenwood et al., 2016; Nagata et al., 2020; Salamango et al., 2019)), Vpr hijacks the CUL4A^DDB1.DCAF1^ E3 ligase complex to target host proteins for ubiquitination, degradation, and cell cycle arrest (reviewed in (Dehart and Planelles, 2008)). Earlier studies demonstrated that induction of cell cycle arrest by Vpr occurs via activation of the DNA damage sensor ataxia telangiectasia and rad3-related (ATR) (Roshal et al., 2003).

Vpr-induced cell cycle arrest is conserved among all *vpr* containing primate lentiviruses and, therefore, likely provides an essential function in maintaining *in vivo* viral fitness (Fletcher et al., 1996; Planelles et al., 1996). Indeed, higher levels of HIV-1 RNA have been measured during the G2 phase of the cell cycle (Gummuluru and Emerman, 1999), and increased integrated provirus was measured following G2/M induced cell cycle arrest (Goh et al., 1998; Yao et al., 1998). These findings suggest that G2/M cell cycle arrest is a phenotype that favors HIV-1 replication. However, a precise understanding of processes that link G2/M cell cycle arrest and virus replication remains elusive.

One of Vpr’s reported functions relating to cell cycle perturbation is transactivation of the viral promoter (Goh et al., 1998; Zhu et al., 2001). A recent report identified the nucleolar protein CCDC137, also known as RaRF as the ubiquitination target of the complex (Zhang and Bieniasz, 2020), and reported transactivation of the viral promoter that was reminiscent of previous observations (Goh et al., 1998; Zhu et al., 2001). The fact that CCDC137 resides within the nucleolus is of particular interest given its capacity to rapidly direct cellular signals to diverse nuclear functions. The nucleolus is a membrane-less subnuclear compartment that, in addition to its canonical role in ribosome biogenesis, coordinates temporally diverse processes such as cell cycle progression, RNA splicing, responses to stress, DNA repair, and apoptosis (reviewed in (Iarovaia et al., 2019). Nucleolar integrity and ribosomal RNA (rRNA) transcription are tightly coupled, with inhibition of rRNA transcription favoring protein translocation from the nucleolus to the nucleoplasm (Iarovaia et al., 2019). CCDC137 was first identified to be a retinoic acid resistance factor (RaRF) that is responsible for sequestering the retinoic acid receptor (RAR) within the nucleolus (Um et al., 2014). CCDC137 was later found to associate with another nuclear receptor, estrogen related receptor alpha (ERRα), and sequester ERRα within the nucleolus (Um et al., 2012). Nucleolar proteins are often redistributed during viral infection, including HIV-1 infection (reviewed in (Salvetti and Greco, 2014)). CCDC137 is antagonized by another viral protein, adenovirus E1A protein, that was shown to disrupt CCDC137 association with RAR, which results in release of RAR into the nucleoplasm and increased transcription of RAR target genes (Um et al., 2014). Recently, Vpr has been shown to target sirtuin 7 (SIRT7), a protein that resides predominantly in the nucleolus (Zhou et al., 2021). However, the consequences of Vpr-mediated degradation of SIRT7 have yet to be determined.

In the present study, we examine the degradation of CCDC137 in the presence of HIV-1 Vpr and its cause-effect relationship with G2/M arrest induction. We confirm that DCAF1 is required for Vpr-induced degradation of CCDC137 and that Vpr-induced G2/M cell cycle arrest is associated with degradation of CCDC137 as previously reported (Zhang and Bieniasz, 2020). However, we observe that degradation of CCDC137 is a general consequence, rather than a trigger, of G2/M arrest. Thus, whether induced by Vpr expression or pharmacologically via CDK1 inhibition, G2-to-M blockage results in degradation of CCDC137.

## Results

### HIV-1 Vpr induces degradation of CCDC137 in a manner that is dependent on DCAF1 and cullin neddylation

We were interested in verifying the ability of HIV-1 Vpr to target CCDC137 for degradation using an orthogonal approach to that previously described (Zhang and Bieniasz, 2020). We and others demonstrated that HIV-1 Vpr mediated cell cycle arrest is dependent on DCAF1 (reviewed in(Dehart and Planelles, 2008)) and reasoned that if HIV-1 mediated degradation of CCDC137 results in cell cycle arrest, degradation of CCDC137 should be dependent on DCAF1. To accomplish this, we transfected 293FT cells with pFIN bicistronic lentiviral vectors (Verrier et al., 2011) expressing either wild-type (WT) Vpr or the Q65R mutant, which fails to bind to DCAF1 (Belzile et al., 2007). We confirm in **Figures 1A** that expression of WT Vpr (lanes 2-6) results in 65-69% degradation of CCDC137, but Q65R (lanes 7-11), results in minimal,12-27% (not significant), degradation of CCDC137, compared to cells transfected with pFIN-GFP (lane 1). The minimal degradation of CCDC137 by Vpr mutant Q65R is likely due to very low affinity between Vpr Q65R and DCAF1. To confirm the dependence of DCAF1 on CCD137 degradation, we next used DCAF1 siRNA to knockdown (KD) DCAF1 protein production. KD of DCAF1 resulted in loss of HIV-1 Vpr mediated degradation of CCDC137 (**Figure 1B**). To confirm the requirement for a cullin-containing ubiquitin E3 ligase complex in the degradation of CCDC137, 293FT cells were treated with the Nedd8 inhibitor MLN4924 after transfection with pFIN-Vpr or pFIN-GFP as a control (**Figures 1C**). In the absence of MLN4924 and in the presence of WT Vpr (lanes 2-4 compared to lane 1), the endogenous CCDC137 protein level was reduced by 57%. However, the endogenous CCDC137 protein level was only reduced by 10% (not significant) in cells treated with MLN4924 (lanes 5-7 compared to lane 1). These results indicate that a cullin-containing ubiquitin ligase is required for CCDC137 degradation because cullin-containing ring ligases require NEDD8 conjugation for their activation (Chiba and Tanaka, 2004; Ramirez et al., 2015; Soucy et al., 2009). Because neddylation of Cul4A was shown to be required for Vpx-induced degradation of sterile alpha motif and histone aspartic domain containing (SAMHD1) protein (Hofmann et al., 2013) and because Vpr increases neddylation of Cul4A in the CUL4A^DDB1.DCAF1^ E3 ligase complex (Hrecka et al., 2007), we asked if neddylation was also required for Vpr-induced cell cycle arrest. To test this idea, we transduced HeLa cells with pFIN-Vpr or pFIN-GFP in the presence or absence of NEDD8 inhibitor MLN4924 (**Figure 1D).** Treatment with MLN4924 partially blocked Vpr mediated cell cycle arrest from (G2+M)/G1=5.4 to 1.9, indicating that neddylation is required.

**Figure 1.**
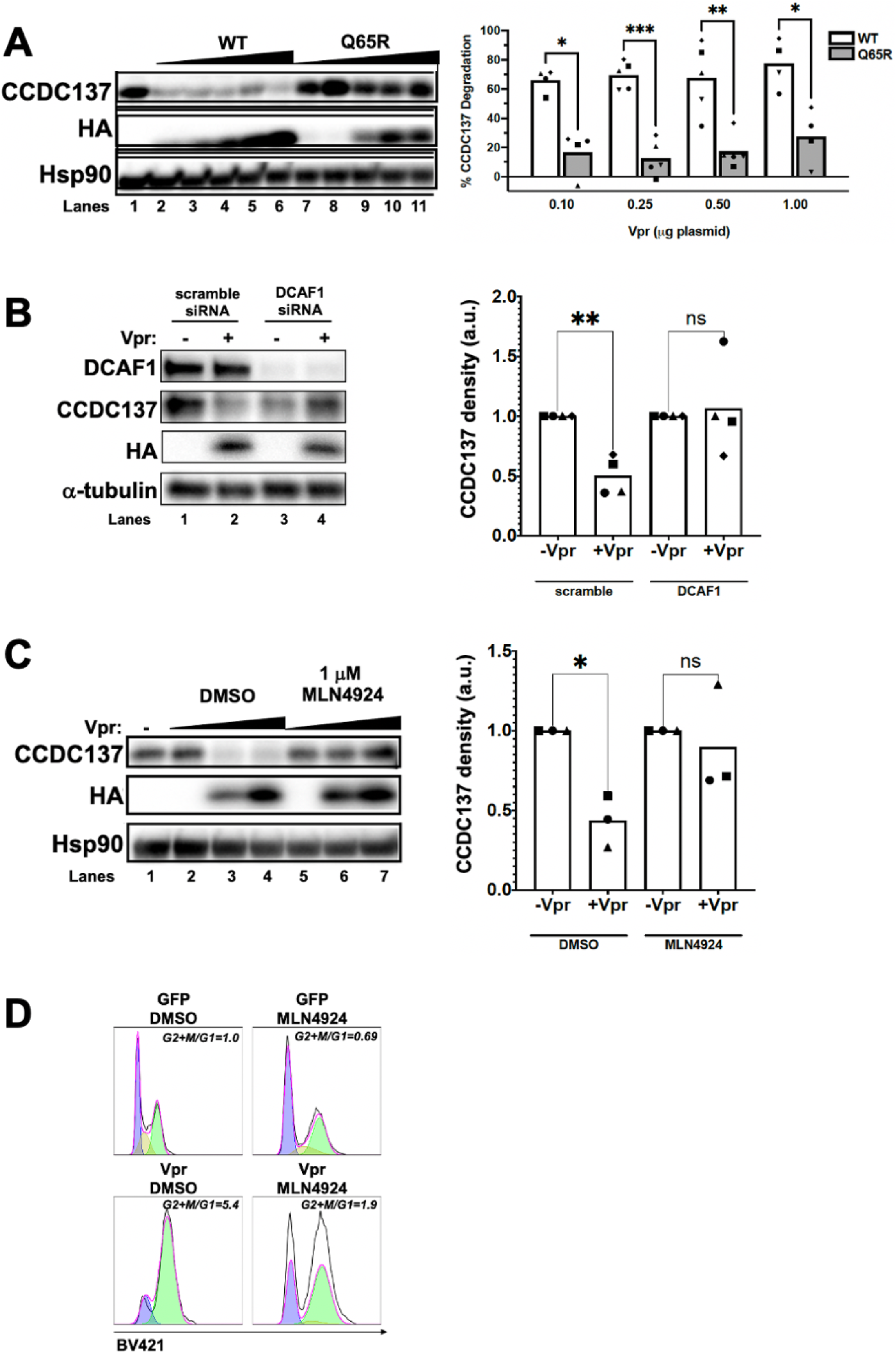
HIV-1 Vpr induces degradation of CCDC137 in a DCAF1 dependent manner. (A) Western blot analysis of endogenous CCDC137, HA tagged HIV-1 VPR, and Hsp90 from 293FT protein lysates 48 hours after transfection with 0.25 μg, 0.50 μg, 0.75 μg, 1 μg, 3 μg pFIN expression constructs: wild-type HIV-1 Vpr (WT), DCAF1 binding mutant Q65R HIV-1 Vpr (Q65R). Gel densiometric quantification from five separate replicate experiments, each replicate displayed as a unique symbol. CCDC137 band densities were normalized to loading control. (B) Western blot analysis of endogenous CCDC137 from 293FT protein lysates 60 hours after transfection with the indicated siRNAs and 48 hours after transfection with pFIN-GFP or pFIN-HAVpr. Gel densiometric quantification from four replicates. CCDC137 band densities were normalized to loading control and then to-Vpr. (C) Western blot analysis of CCDC137, HA tagged Vpr and Hsp90 from 293FT protein lysates 48 hours after transfection with 0 μg, 0.5 μg, 1.0 μg pFIN-Vpr. Cells were treated with DMSO (0.1%) or 1 μM MLN4924 6 hours after transfection. Gel densiometric quantification from four replicates. CCDC137 band densities were normalized to loading control and then to -Vpr. (D) Cell cycle analysis 48 hours after transduction of HeLa cells with the indicated pFIN lentiviruses. Cells were treated with DMSO (0.1%) or 1 μM MLN4924 6 hours after lentiviral transduction. Cells were stained with FxCycle™ Violet DNA dye before analysis by flow cytometry. Cell cycle histograms indicated are for GFP+ transduced cells. P-values were determined using a paired, two-tailed student t-test. . p>0.05 = ns, 0.01<p<0.05 =*, 0.001<p<0.01 = **, 0.0001<p<0.001 = ***, a.u.: arbitrary units

### HIV-1 Vpr cell cycle arrest mutant R80A induces degradation of CCDC137

The HIV-1 Vpr mutant R80A binds to DCAF1 but fails to induce cell cycle arrest because it binds to DCAF1 but presumably fails to recruit the target protein (DeHart et al., 2007). We would expect that VprR80A would fail to degrade CCDC137 and would abolish CCDC137 degradation when co-transfected with wild-type Vpr if CCDC137 is indeed the cell cycle arrest target. However, after transfection with pFIN-VprR80A (lanes 6-10 **Figure 2A** and **Figure 2B**), we observed that degradation of CCDC137 occurred similarly to that observed with WT Vpr (lanes 2-5 **Figure 2A** and **Figure 2B**). This result would indicate that the cell cycle arrest and CCDC137 functions are not interdependent. The mechanism by which Vpr R80A induces degradation of CCDC137 is uncertain.

**Figure 2.**
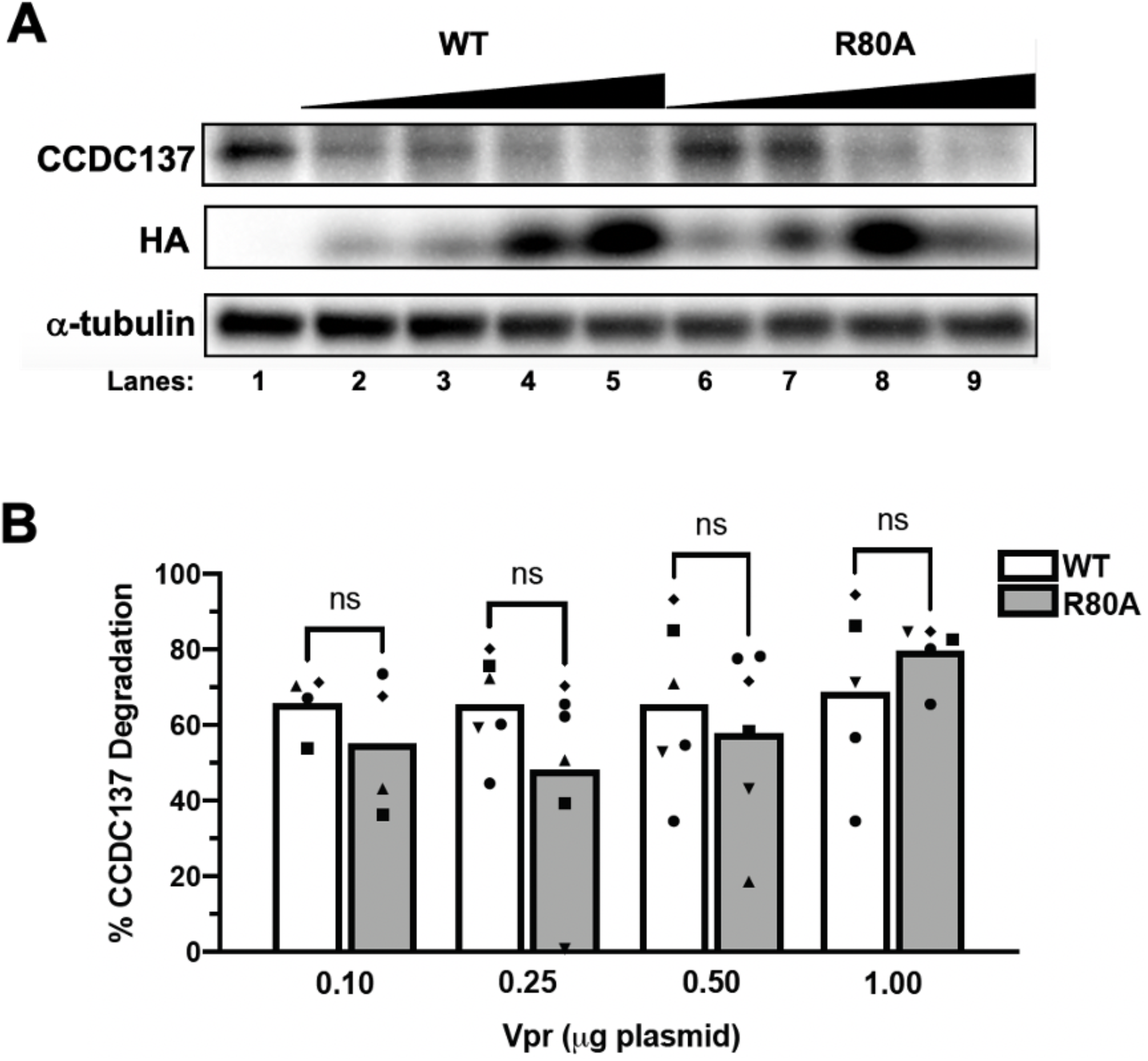
Mutation of Vpr amino acid residue from R to A does not impair binding and degradation of endogenous CCDC137. (A) Western blot analysis of endogenous CCDC137, HA tagged HIV-1 VPR, and a-tubulin from 293FT protein lysates 48 hours after transfection (3 μg/well) with 0 μg, 0.10 μg, 0.25 μg, 0.50 μg, 1.0 μg pFIN expression constructs: wild-type HIV-1 Vpr (WT) or mutant HIV-1 Vpr (R80A). Control plasmid (GFP) was used to achieve a constant plasmid mass. Representative of three experiments. (B) CCDC137 degradation for five replicate experiments of (A). P-values were determined using a paired, two-tailed student t-test. p>0.05 = ns

### Depletion of CCDC137 in HeLa cells using siRNA does not result in G2/M cell cycle arrest

Previous studies demonstrated that depletion of CCDC137 in 293FT or U2OS results in G2/M cell cycle arrest (Zhang and Bieniasz, 2020) and we sought to reproduce this phenotype. We included minichromosome maintenance 10 replication initiation factor (MCM10) as a positive control since MCM10 degradation has been clearly shown to be enhanced by Vpr and to induce cell cycle arrest in multiple reports (Chang et al., 2020; Chattopadhyay and Bielinsky, 2007; Romani et al., 2015). To determine whether CCDC137 depletion results in G2/M cell cycle arrest, HeLa cells were transfected with either a control, non-specific scrambled siRNA or CCDC137 or MCM10 siRNAs. Endogenous levels of CCDC137 and MCM10 were measured by Western blot (**Figure 3A**) and cell cycle analysis was conducted using flow cytometry (**Figure 3B**). Knockdown of MCM10 resulted in partial G2/M cell cycle arrest consistent with previous reports, with the mean (G2+M)/G1 ratio increasing from 0.24 to 0.97 (p=0.001). However, knockdown of CCDC137 did not result in measurable G2/M cell cycle arrest, with mean (G2+M)/G1 ratios of 0.24 and 0.22 (p=0.70) for scrambled siRNA and CCDC137 siRNA treated cells, respectively (**Figure 3B**). Western blot analysis of protein levels for CCDC137 and MCM10 confirms that protein levels were reduced upon transfection with siRNAs (**Figure 3A**). Progression through the G2/M checkpoint requires an active Cyclin A or Cyclin B/cyclin dependent kinase 1 (CDK1) complex (reviewed in (Otto and Sicinski, 2017)). Vpr-induced G2/M cell cycle arrest is associated with inactivating phosphorylation of CDK1 on T14 and Y15 (He et al., 1995; Jowett et al., 1995; Re et al., 1995). We confirm that Vpr induces inactivating phosphorylation of CDK1 upon transfection of 293FT cells with pFIN-HAVpr (**Figure 3C**). The mean pCDK1(Y15) protein band density increased from 0.85 to 1.23 a.u. (arbitrary units), (p=0.041) after transfection of 1 μg pFIN-HAVpr and to 1.47 a.u. (p=0.021) after transfection of 3 μg pFIN-HAVpr. If degradation of CCDC137 resulted in G2/M cell cycle arrest, we would expect that KD of CCDC137 (**Figure 3D**) would result in increased inhibitory phosphorylation of CDK1 on Y15. However, depletion of CCDC137 did not phenocopy Vpr-induced inhibitory phosphorylation of CDK1 (**Figure 3C**). The mean pCDK1(Y15) protein band intensity remained unchanged upon siRNA depletion of CCDC137 relative to non-specific, scrambled siRNA treated cells with mean protein band densities of 1.01 and 0.92 a.u. (arbitrary units), respectively (p=0.39).

**Figure 3.**
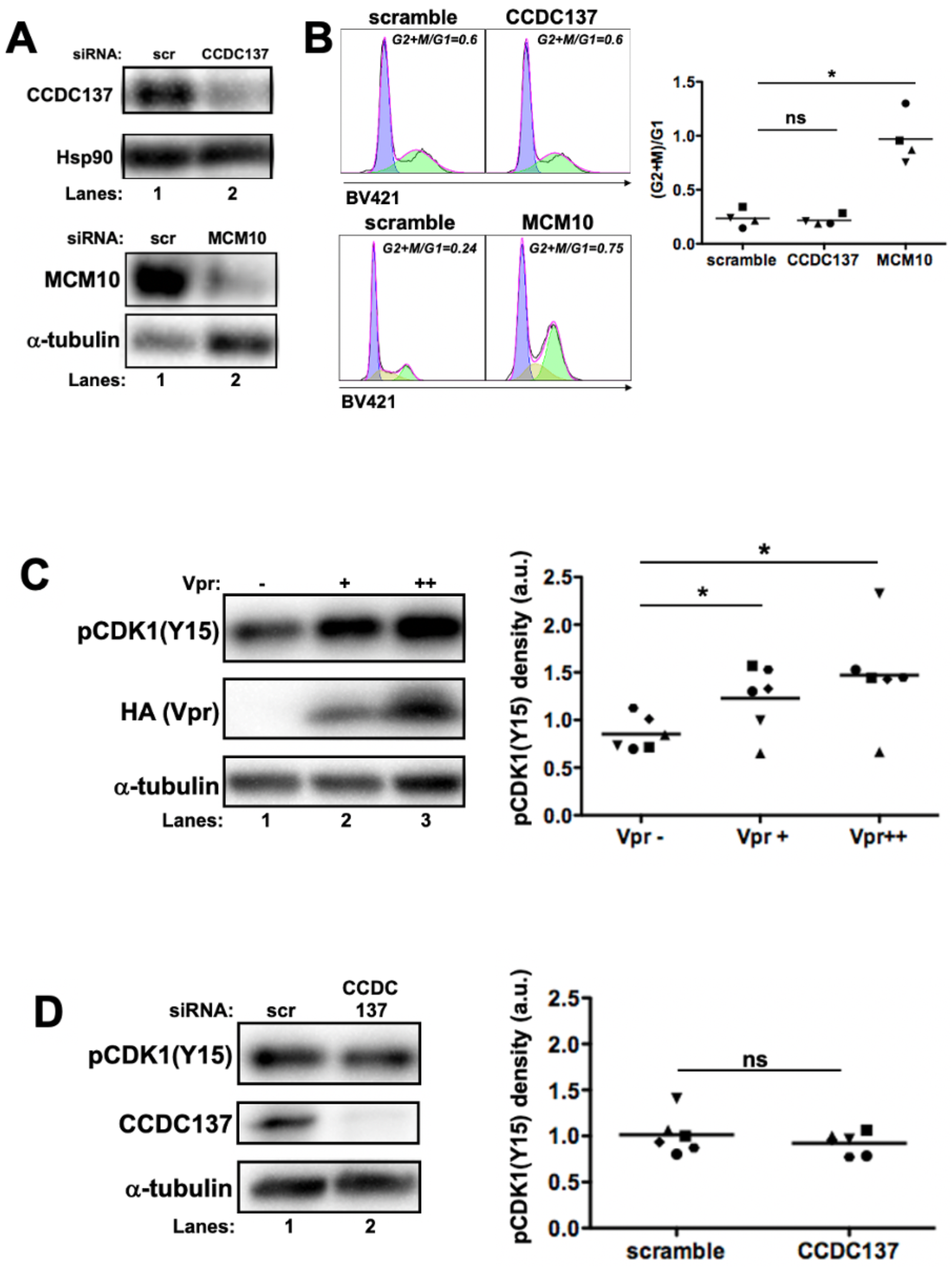
CCDC137 depletion does not result in G2/M cell cycle arrest. (A) Western blot analysis of protein lysates from Hela cell aliquots 48 hours after transfection with siRNAs targeting CCDC137, MCM10 or scrambled. Endogenous CCDC137, MCM10, a-tubulin and Hsp90 protein levels were analyzed. (B) Cell cycle analysis Hela cell sample aliquots from (A) were stained with FxCycle™ violet DNA stain prior to analysis by flow cytometry. Cell cycle histograms indicated are for GFP+ transduced cells. (G2+M)/G1 values for four replicate experiments. Each symbol denotes a separate experiment. (C) Western blot analysis of 293FT transfected with pFIN-GFP (−), 1 μg pFIN-HAVpr (+), or 3 μg HAVpr (++). Endogenous CCDC137, pCDK1(Y15), HA and β-actin protein levels were analyzed. Gel densiometric quantification from six replicates, each replicate displayed as a unique symbol. CCDC137 band densities were normalized to loading control. (D) Western blot analysis of 293FT extracts transfected with scrambled and CCDC137 siRNAs. Endogenous CCDC137, pCDK1(Y15), HA and β-actin protein levels were analyzed. Gel densiometric quantification from six replicates, each replicate displayed as a unique symbol. CCDC137 band densities were normalized to loading control. P-values were determined using a paired, two-tailed student t-test. p>0.05 = ns, 0.01<p<0.05 = *, a.u.: arbitrary units

### Pharmacological induction of G2/M cell cycle arrest results in depletion of CCDC137

Given that knockdown of CCDC137 did not result in G2/M cell cycle arrest, we hypothesized that degradation of CCDC137 may be consequence of G2/M cell cycle arrest. To test this hypothesis, we arrested HeLa cells at the G2/M boundary by incubation with increasing CDK1 inhibitor Ro3306 concentrations and analyzed cell cycle arrest using flow cytometry (**Figure 4A**) and protein levels by Western blot (**Figures 4B** and **4C**). CCDC137 protein levels diminished upon treatment with Ro3306, highly reminiscent of degradation in the presence of Vpr. However, MCM10 protein levels remained unchanged, as expected. Partial G2/M cell cycle arrest was observed with addition of Ro3306 to a concentration of 5 μM and nearly complete cell cycle arrest was observed at 10 μM Ro3306. This result implies that degradation of CCDC137 is dependent on cell cycle arrest.

**Figure 4.**
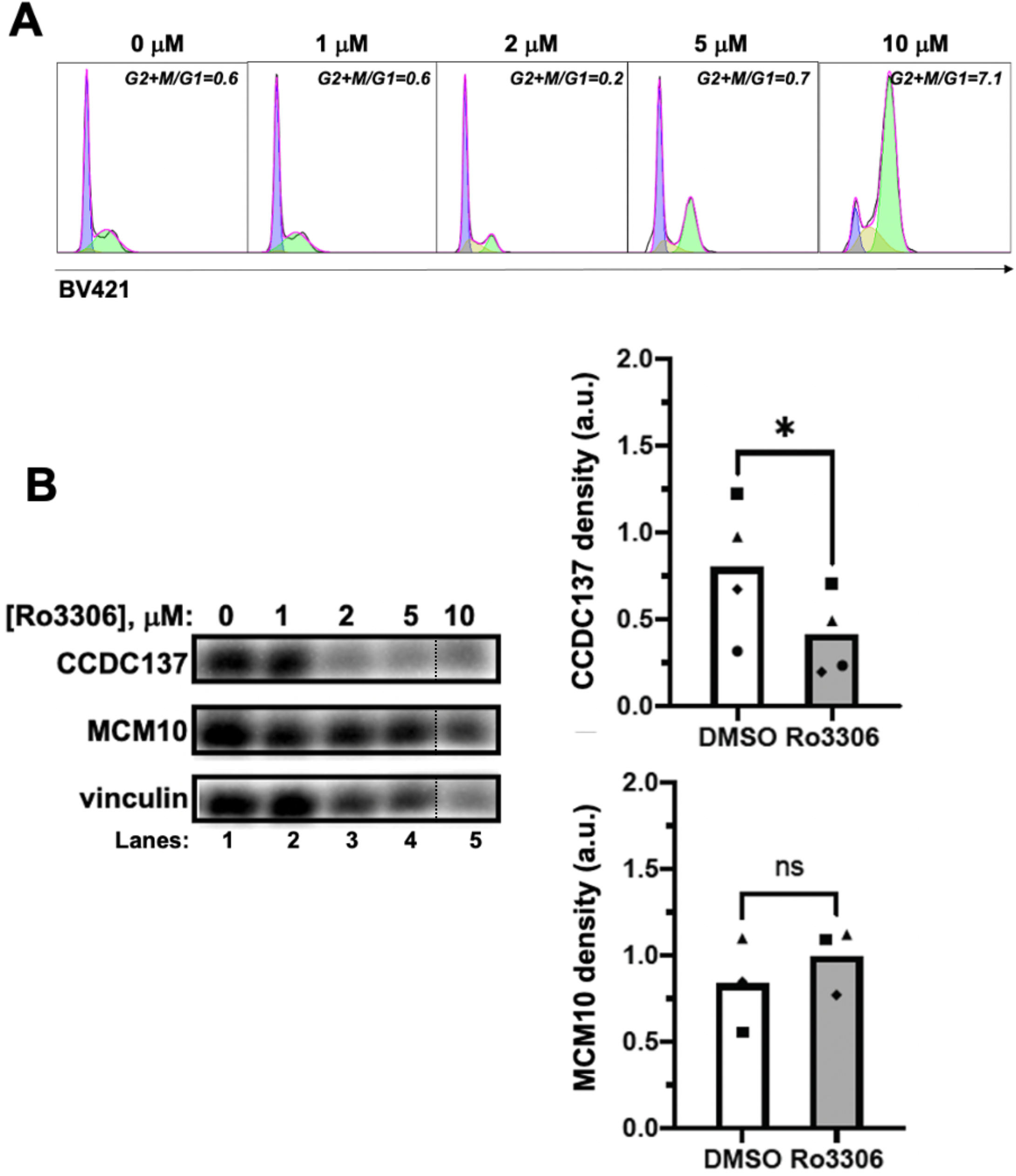
HIV-1 Vpr-induced degradation of CCDC137 is a result of G2/M cell cycle arrest. (A) Hela cells were treated with 0 μM, 1 μM, 2 μM, 5 μM, 10 μM Ro3306. DMSO levels were maintained at 0.1%. After 48 hours, cells were stained with FxCycle™ violet stain prior to analysis by flow cytometry. Cell cycle histograms indicated are for GFP+ transduced cells. (B) Western blot analysis of protein lysates from Hela cell culture aliquots from (A). Endogenous CCDC137, MCM10, vinculin protein levels (loading control) were analyzed. One lane on Western blot image was removed along dotted line. Displayed lanes were exposed and analyzed simultaneously. Gel densiometric quantification from four replicates of (B) Each symbol denotes a separate experiment. CCDC137 band densities were normalized to loading control. P-values were determined using a paired, two-tailed student t-test. p>0.05 = ns, 0.01<p<0.05 = *, a.u.: arbitrary units

## Discussion

Vpr, the only HIV-1 accessory protein known to be encapsidated, has a profound effect on the cellular proteome of the infected cell without requiring *de novo* protein synthesis (Greenwood et al., 2019). Many of these protein abundance changes are dependent on binding to DCAF1, the E3 ligase substrate adaptor, but are not associated with cell cycle arrest (Greenwood et al., 2019). It is therefore compelling to identify proteins that are the ubiquitination targets of the CUL4A^DDB1.DCAF1^ complex resulting in G2/M cell cycle arrest.

A recent report identified the nucleolar protein CCDC137 as the ubiquitination target of the Vpr/CUL4A^DDB1.DCAF1^ complex (Zhang and Bieniasz, 2020) that leads to G2/M cell cycle arrest, and reported transactivation of the viral promoter that was reminiscent of previous observations (Goh et al., 1998; Zhu et al., 2001). In this study, we aimed to confirm and extend the observations by Zhang et al. (Zhang and Bieniasz, 2020).

We report that Vpr mediated degradation of CCDC137 requires the CUL4A^DDB1.DCAF1^ ligase complex. CCDC137 was not significantly destabilized upon siRNA depletion of DCAF1 in the presence of Vpr or by mutation of residue Q65 to arginine. This result agrees with previous proteomic results that identified CCDC137 as a protein potently depleted by WT but not Q65R Vpr. We additionally demonstrated that the Vpr cell cycle arrest null mutant R80A fails to suppress degradation of CCDC137 a finding that also agrees with previous proteomic results (Greenwood et al., 2019), in which depletion of CCDC137 in the presence of cell cycle null mutant S79A was similar to that observed for WT Vpr. We found that siRNA depletion of CCDC137 did not induce cell cycle arrest whereas depleting MCM10 by the same method did lead to G2/M cell cycle arrest, as expected (Chang et al., 2020; Chattopadhyay and Bielinsky, 2007; Romani et al., 2015). Consistent with this result, CCDC137 depletion did not induce inhibitory phosphorylation of CDK1 at Y15, which is a requirement for arrest at the G2/M boundary (Jin et al., 1996). On the other hand, pharmacologically induced cell cycle arrest at the G2/M boundary, enforced through CDK1 inhibition, did lead to CCDC137 destabilization. These results indicate that G2/M cell cycle arrest leads to CCDC137 destruction, but not vice versa.

The use of cell-cycle arrest null mutants has been used extensively to decouple Vpr’s protein degradation and cell cycle arrest functions. For example, MUS81(DePaula-Silva et al., 2015), helicase-like transcription factor (HLTF) (Lahouassa et al., 2016), tet methylcytosine dioxygenase 2 (TET2) (Lv et al., 2018), and SIRT7(Zhou et al., 2021) are targeted for depletion by WT Vpr and by cell-cycle arrest null mutants, such as R80A, indicating that degradation of these cellular targets are not associated with cell-cycle arrest. However, steps leading the degradation of CCDC137 by cell cycle null R80A Vpr mutant are unclear.

CDK1 is a key factor in the tight coordination between nucleolar integrity and cell cycle progression (Hayashi et al., 2018). CDK1 inactivation at the onset of mitosis suppresses rRNA transcription leading to disassembly of the nucleolus and translocation of numerous proteins to the nucleoplasm (Hayashi et al., 2018). At the G2/M transition, CDK1 is activated, rRNA transcription resumes, and the nucleolus reassembles (Hayashi et al., 2018). Given the finding that Vpr expression is associated with degradation of nucleolar proteins CCDC137 (Zhang and Bieniasz, 2020) and SIRT7 (Zhou et al., 2021), it is plausible that Vpr cell cycle arrest may favor viral replication and transactivation through delayed assembly of the nucleolus as a consequence of CDK1 inhibitory phosphorylation. By analogy to adenovirus E1A protein antagonism of CCDC137 to enhance RAR transactivation, it is also possible that Vpr binding and degradation of CCDC137 displaces a protein, such as a transcription factor, that is necessary for enhanced viral transcription and replication. In addition, despite vigorous efforts to define cellular factors targeted by Vpr required for initiation of G2/M cell cycle arrest, further investigations are warranted.

## Materials and Methods

### Materials

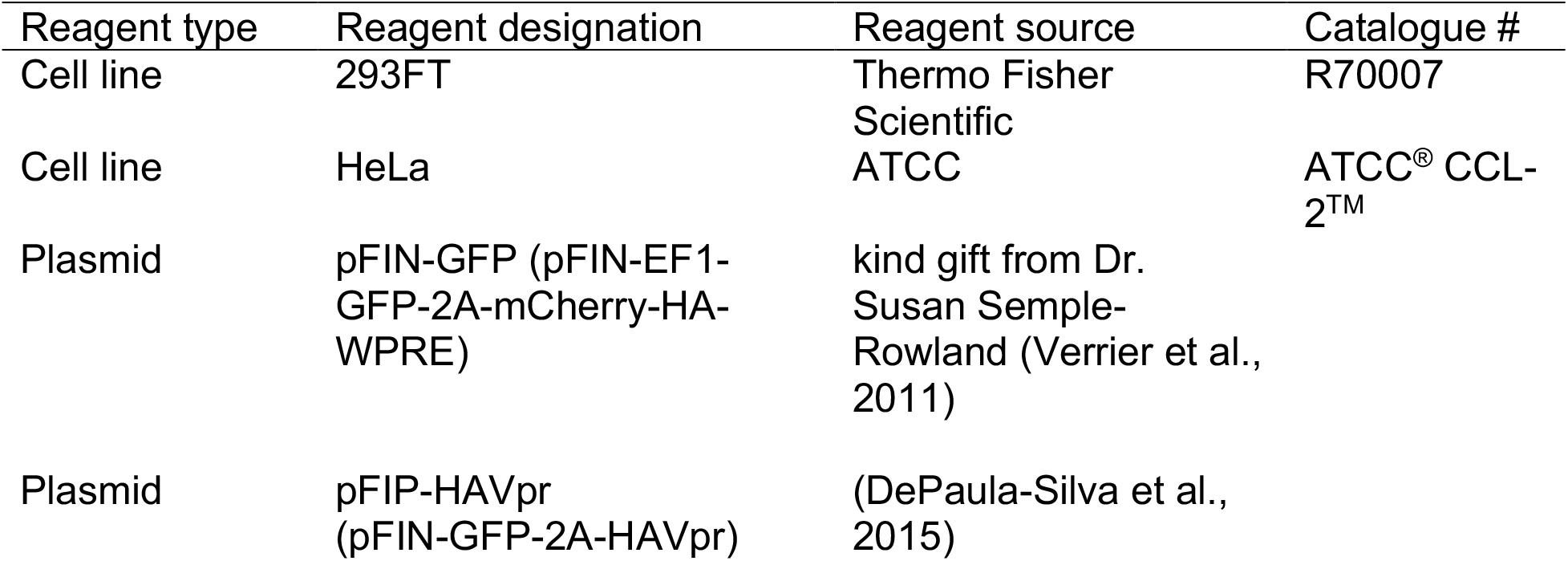

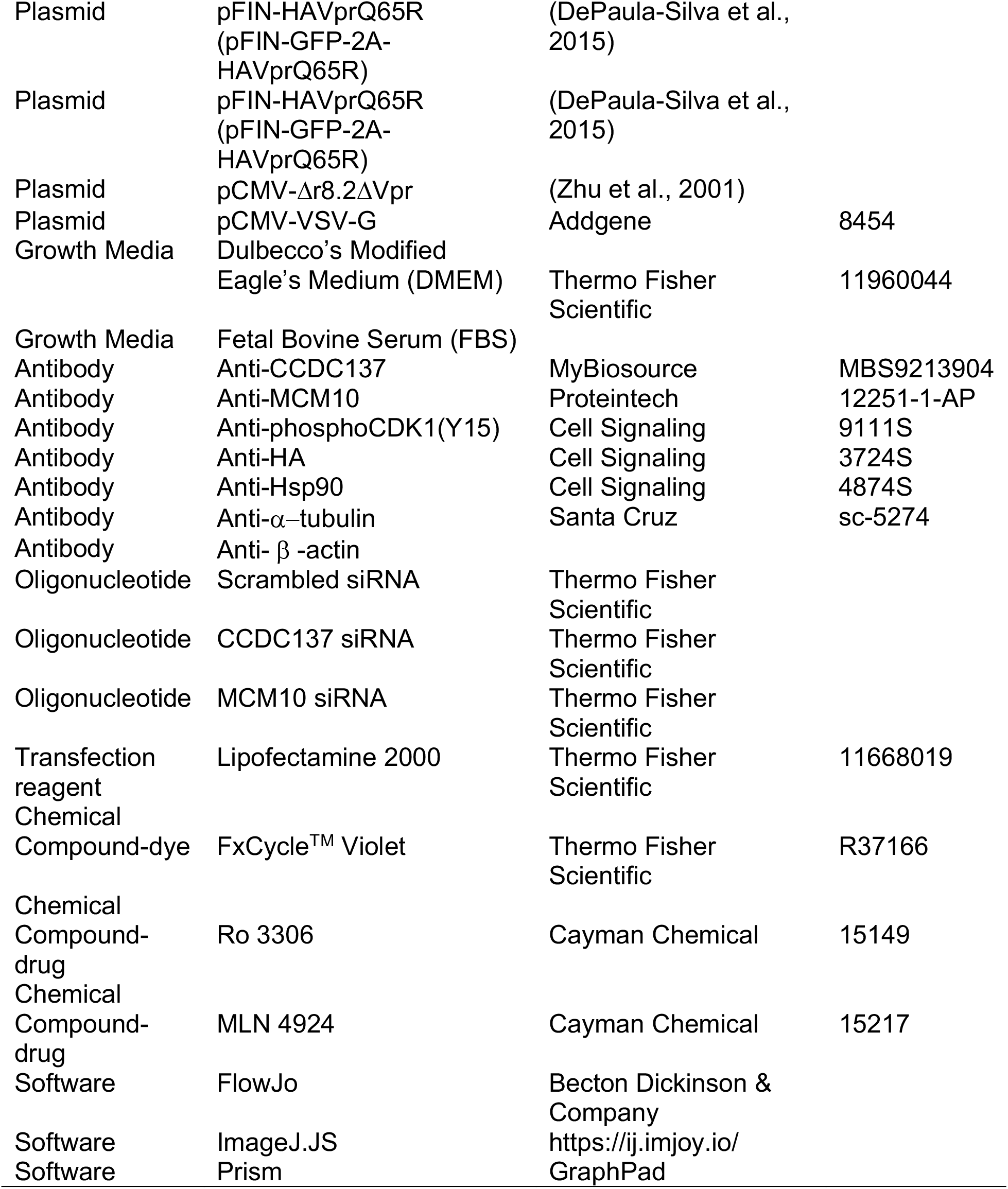

#### Cell lines and transfection

293FT cells and HeLa cells were cultured in Dulbecco’s Modified Eagle’s Medium supplemented with 10% FBS. 293FT and HeLa cell were transfected with siRNAs following the manufacturers protocol. 293FT cells in 6-well plates were transfected with expression plasmids using calcium phosphate as follows. Briefly, 3 μg of pFIN plasmid or plasmids were diluted to a volume of 65 μL in ddH2O. 7.5 μL of 2.5 M CaCl2 and 72.5 μL were added, tubes vortexed and incubated for 1 minute and added to wells dropwise followed by addition of 1.4 μL of 100 mM chloroquine.

#### Virus production and transduction

Lentiviral vectors were produced in 293FT cells. Briefly 12.5 μg transfer plasmid (pCMV-Δr8.2Δvpr), 12.5 μg pFIN lentiviral plasmid, and 5 μg pCMV-VSV-G envelope expressing plasmids were co-transfected into 293 cells using calcium phosphate as previously described. Supernatants were collected after 48 hours and lentiviral vectors concentrated by ultracentrifugation at 25,000 RPM for 2 hours at 4 °C. HeLa cells in 6-well plates were transduced with lentiviral vectors by incubation in the presence of 0.01 mg/mL polybrene for 6 hours.

#### Immunoblotting

Cells were washed in PBS and lysed in NTEN buffer supplemented with phosphatase and protease inhibitors (Roche). Protein concentrations were determined using a BCA assay (Pierce). Protein lysates were diluted to 25 μg in 15 μL in Lamelli buffer and boiled for 10 minutes. Denatured proteins were subjected to SDS-PAGE on 4-15% Criterion™ Tris-Glycine gels (Biorad). Proteins were transferred to Nitrocellulose membranes (0.45 μm, Thermo Fisher Scientific). Membranes were blotted with primary antibodies, indicated in materials table, followed by blotting with HRP-conjugated secondary antibodies, then exposed using chemiluminescence (Pierce ECL, Thermo Fisher Scientific). Western blot bands were quantified using ImageJ.JS.

#### Cell Cycle analysis

Cell cycle analysis was conducted as follows. Briefly, HeLa cells were detached using trypsin, washed with PBS, then fixed with 0.25% paraformaldehyde. Cells were resuspended in 200 μL PBS+3% FBS + 6 μL of FxCycleTM Violet Ready Flow™ reagent and incubated at room temperature for 30 minutes. Cells were subjected to flow cytometry and cell cycle analyzed using FlowJo.

## Disclosure of potential conflicts of interest

The authors declare no conflicts of interest.

## Funding

5R01AI143567-02 Planelles/Coiras (PIs); 11/07/2018-10/31/2023 NIH/NIAID

**Tyrosine Kinase Inhibition: The New Front in HIV Cure Efforts**

## Author contributions

LM designed and conducted experiments and prepared manuscript. ABDS cloned pFIN Vpr expression plasmids, consulted on experimental design, and edited manuscript. VP initiated the project, consulted on experimental design, and revised manuscript.

## Notes

### Competing Interest Statement

The authors have declared no competing interest.

